# Extract from Polygala fallax Hemsl. Protects Kidneys in db/db Mice by Inhibiting the TLR4/MyD88/NF-*κ*B Signaling Pathway

**DOI:** 10.1101/2023.08.08.552432

**Authors:** Yukun Bao, Zeyue Wang, Qing Xu, Lixin Wang, Yi Wen, Peng Deng, Qin Xu

## Abstract

Diabetic nephropathy (DN) is a chronic kidney disease caused by the loss of renal function. The extract of Polygala fallax Hemsl (EPF) possesses anti-inflammatory a nd other pharmacological effects. Objective: To investigate the effect and potential mechanism of EPF in the treatment of diabetic nephropathy-associated inflammation. Materials and methods: Db/db mice were administered varying doses of EPF (15, 30, 60 mg/kg), after which the kidney organ index and glucose tolerance were calcu lated. Urine microalbumin was detected in urine collected over 24 hours. Serum FBG, Cr, and BUN levels were measured, and H&E and PAS staining were used to observe pathological changes in the kidney. The expression of TLR4, MyD88, NF-*κ*B, and MMP −9 in kidney tissue was measured using immunohistochemistry, quantitative real-tim e PCR, and western blotting. Additionally, the expression of TNF-*α*, MCP-1, IL-6, IL-18, and IL-1*β* inflammatory factors in the serum was measured by ELISA. Results : EPF significantly decreased the renal organ index and ameliorated glucose intolerance symptoms in db/db mice, reduced 24-hour mALB, FBG, Cr, and BUN serum levels, and mitigated renal pathological changes. Moreover, EPF significantly inhibited th e expression of TLR4, MyD88, NF-*κ*B, MMP-9, and related inflammatory factors TNF-*α*, MCP-1, IL-6, IL-18, and IL-1*β* in kidney tissue. Discussion and conclusions: E PF from P. fallax exhibits low toxicity and is safe for use. For the first time, it was discovered that EPF might reduce renal inflammation by inhibiting the TLR4/M yD88/NF-*κ*B signaling pathway in vivo, thereby protecting the kidneys of db/db mic e from damage.

## Introduction

Diabetes mellitus (DM) is a group of diseases characterized by elevated plasma glu cose levels. With the sharp increase in the number of elderly people in many count ries, the prevalence of diabetes has also risen year by year^1^. DM has become a major chronic non-communicable disease posing a serious threat to health, alongside tumors and cardiovascular and cerebrovascular diseases. Epidemiological studies hav e shown that in 2015, the number of people with diabetes worldwide surpassed the total population of the United States, reaching 415 million^2^. As of 2019, approximately 463 million people worldwide have diabetes^3^.

Significant progress has been made in the clinical treatment of diabetic patients through the application of insulin. However, with the extension of life expectancy for diabetic patients, chronic complications have become the main concern. According to statistics, 20–50% of diabetic patients will develop diabetic nephropathy (D N)^4^. In China, approximately 21.3% of diabetic patients will eventually develop DN^5^. Clinically, the entire course of DN is primarily accompanied by a continuous increase in proteinuria and a severe decrease in the glomerular filtration rate (GFR)^6^. At present, there is no effective treatment for DN. Despite active anti-diabetic treatments, blood pressure reduction, weight control, and the administration of angiotensin-converting enzyme inhibitors (ACEI) or angiotensin receptor blockers (ARBs), anticoagulation, lipid regulation, and other comprehensive treatments, the incidence of end-stage renal failure in patients with DN remains high^7^. Given the significant threat posed by DN to the health of diabetic patients, it is urgent to explore new effective treatment strategies for DN.

The innate immune system serves as the first line of defense against pathogenic microorganisms and stimulates the innate immune response through pattern recognition receptors (PRRs). To date, several PRRs have been identified, including toll-like receptors (TLRs) and nucleotide-binding oligomerization domain (NOD)-like receptor s (NLRs)^8^. TLRs are the most extensively studied, with toll-like receptor 4 (TLR4) being the first discovered and well-known among all human TLRs. TLR4 is widely expressed in renal intrinsic cells, such as mesangial cells, renal tubular epithelial cells, and podocytes^9^.

In recent years, researchers have found that inflammation plays an important role in the occurrence and development of DN, and DN induces various factors to promote inflammation^10^. The TLR4/NF-*κ*B pathway is an important regulatory pathway for inf lammatory responses^11^. TLR4 is activated by binding with endogenous ligands released from immune and kidney cells, initiating a downstream signaling cascade through MyD88-dependent and -independent pathways, ultimately leading to the activation of Nuclear factor kappa B (NF-*κ*B)^12^. Activated NF-*κ*B is then transferred to the nuc leus, inducing the transcription and translation of related inflammatory mediator genes, resulting in an increased release of pro-inflammatory cytokines and chemoki nes^13^. Therefore, effectively decreasing renal inflammation is key to reducing DN damage^14^.

False yellowflower milkwort/yellow ginseng (Polygala fallax Hemsl.; Order: Polygal ales, Family: Polygalaceae) is a commonly used folk medicine in Guangxi. Its main chemical components are flavonoids and saponins, which have notable effects on lowering blood lipids and decreasing myocardial ischemia, as well as anti-fatigue, an ti-oxidation, anti-aging, anti-inflammatory, and anti-viral properties^15,16^. The extract of P. fallax (EPF) is derived from its root and stem, with its bioactive component being the parent nucleus structure of pentacyclic triterpenoids. Previous studies have shown that P. fallax can decrease the expression of Tumor necrosis factor-*α* (TNF-*α*) and Interleukin-6 (IL-6) inflammatory factors, thereby halting the development of glomerulonephritis^3^. However, the mechanism by which EPF mitigates diabetic nephropathy remains unclear. Therefore, this study aims to explore the renal protection offered by EPF on DN and its related mechanisms, providing a strong experimental basis for the clinical treatment of DN.

## 2. Materials and Methods

### 2.1 Animals

Specific pathogen-free (SPF)-grade male C57BL/KsJ/db-/-mice (five weeks old) were acquired from GemPharmatech Co., Ltd. (Nanjing, China, license number: SCXK2018-00 08). SPF-grade Kunming (KM) mice (five weeks old) were sourced from Hunan SJA Laboratory Animal Co., Ltd. (Changsha, China, license number: SCXK (Hunan) 2016-0002). The mice were provided with food and water ad libitum and housed at 20–26°C (temperature variation not exceeding 4°C) with 40%–70% humidity. All experiments were approved by the laboratory animal ethics committee and conducted in accordance wit h the guidelines for the care and use of laboratory animals.

### 2.2 Drugs and Reagents

EPF was supplied by Nanjing Puyi Biological Products Co., Ltd. (Nanjing, Jiangsu); Report Number: Py-395–201031. The absorbance of EPF was measured using tenuifolin as the standard, and the mass concentration of tenuifolin in the test solution was determined from the standard curve. EPF was uniformly suspended in 0.5% sodium carboxymethyl cellulose (CMC-Na) for intragastric administration. Gliquidone tablets (GLI) were obtained from Beijing Wanhui Double-Crane Pharmaceutical Co., Ltd. (Bei jing, China).

### 2.3 Experimental Design

The db/db mice were randomly assigned to five groups, with six mice per group, and six db/m mice were placed in the control group. After one week of acclimation, different concentrations of EPF or GLI were administered daily: positive group (10 mg/kg gliquidone), EPF low-dose group (15 mg/kg), EPF medium-dose group (30 mg/kg), and EPF high-dose group (60 mg/kg). The blank and model groups received the same d ose of 0.5% CMC-Na for eight weeks. A portable blood glucose meter (Accu-Chek Active Blood Glucose Meter, Roche, Cork, Ireland) was employed to assess the fasting blood glucose (FBG) levels of the mice, and their weights were recorded weekly.

### 2.4 Oral Glucose Tolerance Test (OGTT)

After the final administration, the mice were fasted for 12 hours, and blood glucose was measured using tail-tip blood sampling, which was recorded as 0-hour blood glucose. The mice were then given 1 g/kg glucose via gavage, and their blood glucose levels were measured at 0.5, 1, and 2 hours after administration. The following formula was used to calculate the glucose area under the curve (AUC): AUC = (0 h blood glucose + 0.5 h blood glucose) × 0.25 + (0.5 h blood glucose + 1 h blood glucose) × 0.25 + (1 h blood glucose + 2 h blood glucose) × 0.5

### 2.5 Kidney/Body Index Calculation

After glucose administration, the mice were fasted for 12 hours and then weighed. Following euthanization, their kidneys were removed, rinsed with normal saline, an d dried with filter paper to remove any residual liquid. The kidneys were then weighed. The formula used to calculate the renal organ index is as follows: renal organ index (%) = left kidney weight (g)/body weight (g) × 100.

### 2.6 Determination of Urine Microalbumin (mALB)

In the ninth week after the end of treatment, all mice were placed in metabolic cages with free access to food and water. Their urine was collected for 24 hours, an d urine output was recorded. After the collected urine had been left undisturbed f or 3 hours, it was centrifuged at 3500 rpm for 10 minutes. The supernatant was col lected, and the level of microalbumin in the urine was detected using an automatic biochemical analyzer (Cobas 8000 modular analyzer series, Roche Diagnostics).

### 2.7 Determination of Serum Parameters

After urine collection, the mice were fasted overnight. The unilateral eyeball of each mouse was carefully removed, and blood was collected. After standing for 3 hours, the blood was centrifuged (3500 rpm, 10 minutes, 4°C) to separate the serum. An automatic biochemical analyzer was employed to measure serum creatinine (Scr) a nd urea nitrogen (BUN) levels (Cobas 8000 modular analyzer series, Roche Diagnostics). ELISA kits (Solarbio, Beijing, China) were used to measure the expression lev els of TNF-*α*, Interleukin-18 (IL-18), Interleukin-1*β* (IL-1*β*), Monocyte Chemotac tic Protein-1 (MCP-1), and IL-6 in mouse serum.

### 2.8 Histopathology and Histochemistry

After euthanizing the mice, their eyeballs were removed, and blood was collected. The mice were then immediately dissected, and the middle third of the right kidney was excised and fixed in 10% neutral formalin for 48 hours. The samples were then embedded in paraffin and sectioned to a thickness of 4 μm for histological examination and immunohistochemical (IHC) analysis. For histological examination, hematoxylin and eosin (H&E) and periodic acid-Schiff (PAS) staining were employed to observe kidney tissue sections (Olympus, BX53, Japan) under a microscope, and images were recorded.

For IHC analysis, after paraffin embedding, 4-μm-thick kidney tissue sections underwent antigen retrieval. Endogenous catalase activity was blocked, and the sections were incubated with TLR4 (Solarbio, Beijing, China) (1:200), NF-*κ*B (Solarbio, Beijing, China) (1:250), MyD88 (Solarbio, Beijing, China) (1:50), and MMP-9 (Solar bio, Beijing, China) (1:200) antibodies overnight at 4°C. The next day, the sections were washed with phosphate-buffered saline (PBS, 5 minutes × 3) and incubated with goat anti-rabbit IgG horseradish peroxidase-conjugated secondary antibody (Solarbio, Beijing, China) (1:500) at 37°C for 50 minutes. After diaminobenzidine (D AB) staining, the expression of each protein was observed with an optical microscope (Olympus, BX53, Japan). The results were carefully analyzed using ImageJ software, and the relative expression was represented by the ratio of the integral optical density (IOD) of the positive area to its area (Area).

### 2.9 Quantitative Reverse Transcription PCR (RT-PCR)

Total RNA from kidney tissue was extracted using TRIzol reagent (Tiangen, Beijing, China). A NanoDrop One spectrophotometer was employed to measure the total mRNA concentration (Thermo Fisher Scientific, USA). The ReverTra Ace-cDNA synthesis kit (Promega, USA) was used to synthesize cDNA from 2 μg of total RNA. Gene expression analysis was performed using an Applied Biosystems™ 7500 Fast Real-Time PCR System (Thermo Fisher Scientific, USA). The primer sequences used for gene expression ana lysis are shown in Table 1. The ABI 7500 Fast v2.0.1 system employed the 2-ΔΔCt method to analyze mRNA fold changes. Primer sequences used for gene expression ana lysis are provided in Table 1.

**Table 1.**
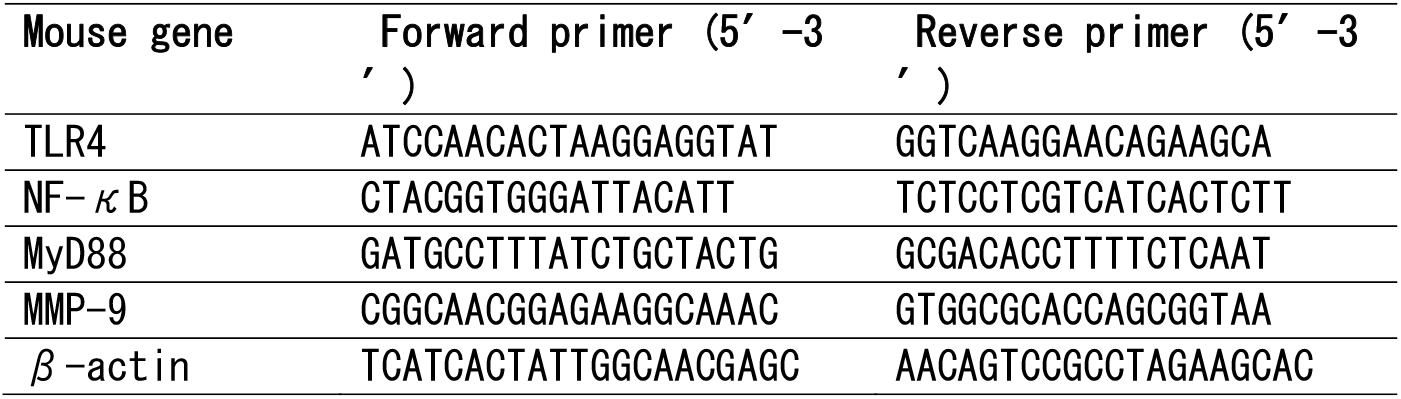
Sequences of primers.

### 2.10 Western Blotting (WB)

Total protein was extracted from kidney tissue using prepared radioimmunoprecipitation assay (RIPA) lysate. After quantification with a bicinchoninic acid (BCA) kit, the total protein was separated by sodium dodecyl sulfate-polyacrylamide gel electrophoresis and transferred to a polyvinylidene difluoride (PVDF) membrane. The P VDF membrane was incubated with 5% skim milk in a sealed container at room tempera ture for 2 hours. Then, the corresponding primary antibodies were added, including TLR4 (CST, Boston, USA) (1:1400), MyD88 (CST, Boston, USA) (1:1400), NF-*κ*B p65 (CST, Boston, USA) (1:1400), and MMP-9 (CST, Boston, USA) (1:1000), and the membrane was incubated overnight at 4°C. *β*-actin (1:5000) was used as an internal reference. The membrane was subsequently incubated with an appropriate secondary antibody (1:10000) in a shaking flask at 37°C for 2 hours. Finally, enhanced chemiluminescence (ECL) reagent (Biosharp, Shanghai, China) was added, and the membrane was exposed in a dark room. An imaging system was employed to quantify the expression of each protein (Bio-Rad, USA).

### 2.11 Acute Toxicity Test in KM Mice

Twelve KM mice (six females and six males) were randomly divided (n = 6) into a blank control group and an administration group, which received 300 times the EPF of the higher dose group. After one week of acclimation, the mice were fasted for 8 hours before the experiment. The administration group was given a single dose of 1. 8 g/kg, and the control group received the same dose of 0.5% CMC-Na. Toxic reactions in the mice were observed and immediately recorded, and the activity of the mic e was closely monitored within 4 hours after administration. After 14 days of continuous observation of the mice’s survival status, the mice were fasted for 8 hours, euthanized, and blood was collected from the eyeballs to measure their albumin (ALB), total protein (TP), serum creatinine, urea nitrogen, alanine aminotransferase (ALT), and aspartate aminotransferase (AST). The mice were dissected to quickly extract the heart, liver, kidney, and spleen, which were subsequently weighed to calculate the aforementioned organ index.

### 2.12 Statistical Methods

GraphPad Prism 5.0 was used for data analysis, and the results are expressed as th e mean ± standard error of the mean (mean ± S.E.M). Comparisons were made using one-way ANOVA followed by Tukey’s test. A P-value < 0.05 was considered statistica lly significant.

## Results

### 3.1 Analysis Results of EPF

EPF was extracted from the roots and stems of P. fallax. The absorbance of EPF and the reference substance were measured at 580 nm, using the solution without the reference substance as a blank to obtain the standard curve. The dosage was 19.2 kg, the weight of the extracted sample was 106.8 g, and the yield was 0.56%, with the content of EPF saponin being 63.3%.

### 3.2 EPF Ameliorated Glucose Intolerance Symptoms in db/db Mice

Glucose tolerance characterizes the body’s ability to regulate blood glucose concentration. After oral consumption of a certain amount of glucose, blood glucose lev els were measured at regular intervals. This experiment helps to understand pancreatic islet *β*-cell function and the body’s ability to regulate blood glucose concentration. After oral glucose administration, the blood glucose levels of db/db and db/m mice significantly increased, reaching a maximum at 0.5 h. Subsequently, the blood glucose levels in each group decreased to varying degrees. Compared with unt reated db/db mice, the EPF high-, medium-, and low-dose groups, and the GLI group, exhibited a significant decrease in blood glucose after 2 h. The blood glucose level and the glucose area under the curve (AUC) were significantly lower than those of the untreated group (P < 0.05) (Fig. 1A-B). These data indicate that EPF can significantly ameliorate glucose intolerance in db/db mice.

**Figure 1.**
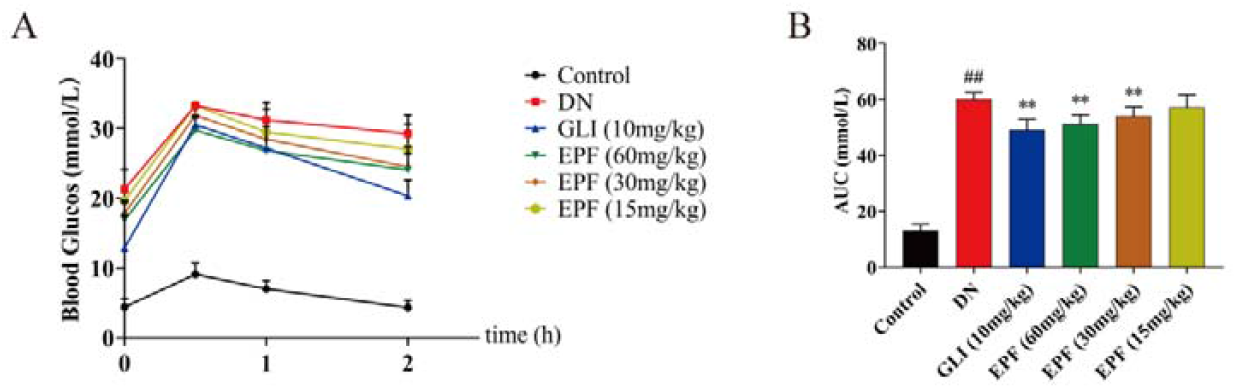
The effect of EPF on OGTT of db/db mice (n=6). (A) OGTT line chart (B) AUC area chart. All data are expressed as mean ± S.E.M. ^##^*P*<0.01 compared with the control group; \**P*<0.05, \*\**P*<0.01 compared with the DN group

### 3.3 EPF Normalized Biochemical Parameters of db/db Mice and Decreased Kidney Damage

Following EPF intervention, compared with the treatment group, the DN group mice began to lose significant amounts of weight, and gradually became drowsy, slow to respond, slow to act, and exhibited symptoms such as diabetic polydipsia, polyuria, and weight loss. There was a significant increase in FBG, and the corresponding symptoms in the treatment group were effectively alleviated. Moreover, the renal organ coefficients of the EPF group and GLI group were significantly lower than those of the DN group, indicating that EPF and GLI effectively reduced renal tissue hypertrophy (Fig. 2A-C). In addition, the 24 h urine microalbumin, blood creatinine, a nd urea nitrogen levels of the DN group were significantly higher than those of th e control group. Under the treatment of EPF and GLI, compared with the DN group, these indices were significantly reduced (Fig. 2D-F). Concurrently, it was found th at compared with the DN group, the expression of TNF-*α*, MCP-1, IL-6, IL-18, and I L-1*β* in the serum was significantly decreased by EPF and GLI in db/db mice (Fig. 3A-E). Therefore, it was concluded that EPF was effective in ameliorating diabetic nephropathy and decreasing inflammation.

**Figure 2.**
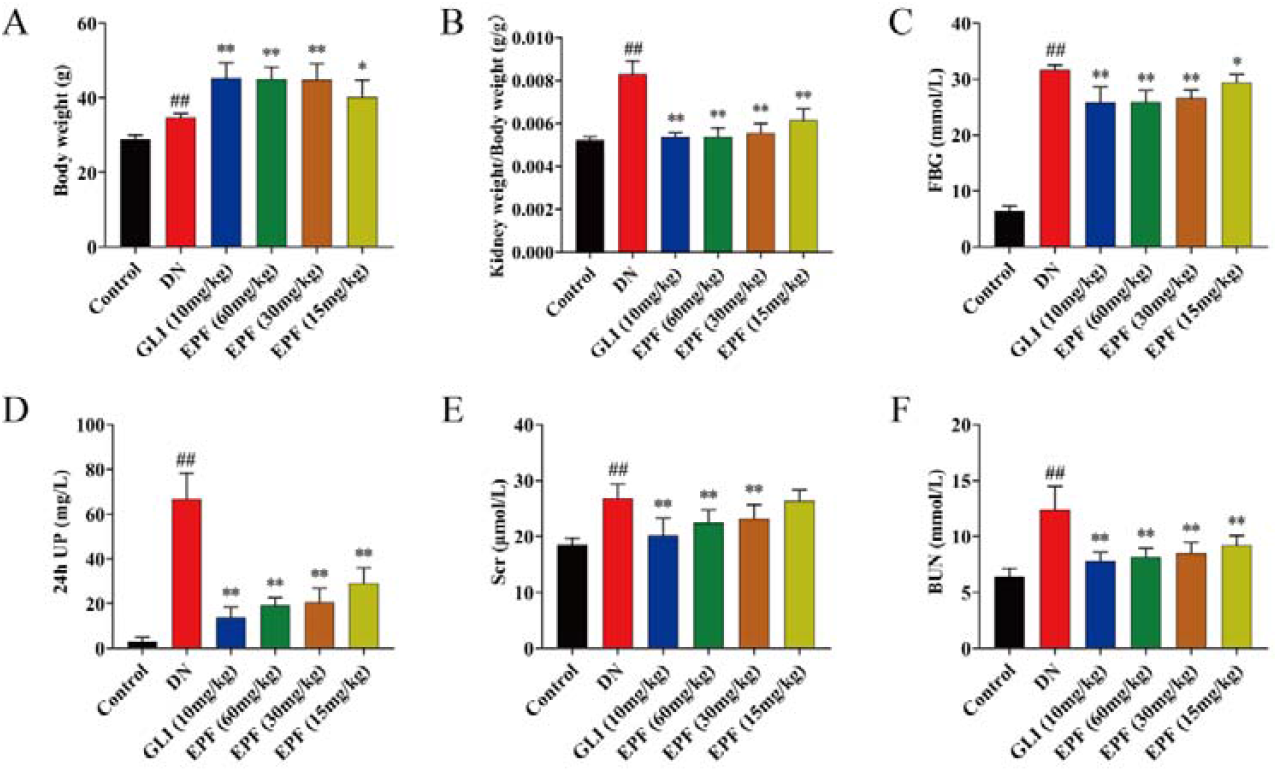
The effect of EPF on the metabolic parameters of db/db mice (n=6). (A) Body weight. (B) Kidney weight/body weight (C) FBG (D) 24h UP (E) Scr (F) BUN. All data are expressed as mea n ± S.E.M. ^##^*P*<0.01 compared with the control group; \**P*<0.05, \*\**P*<0.01 compared with the DN group

**Figure 3.**
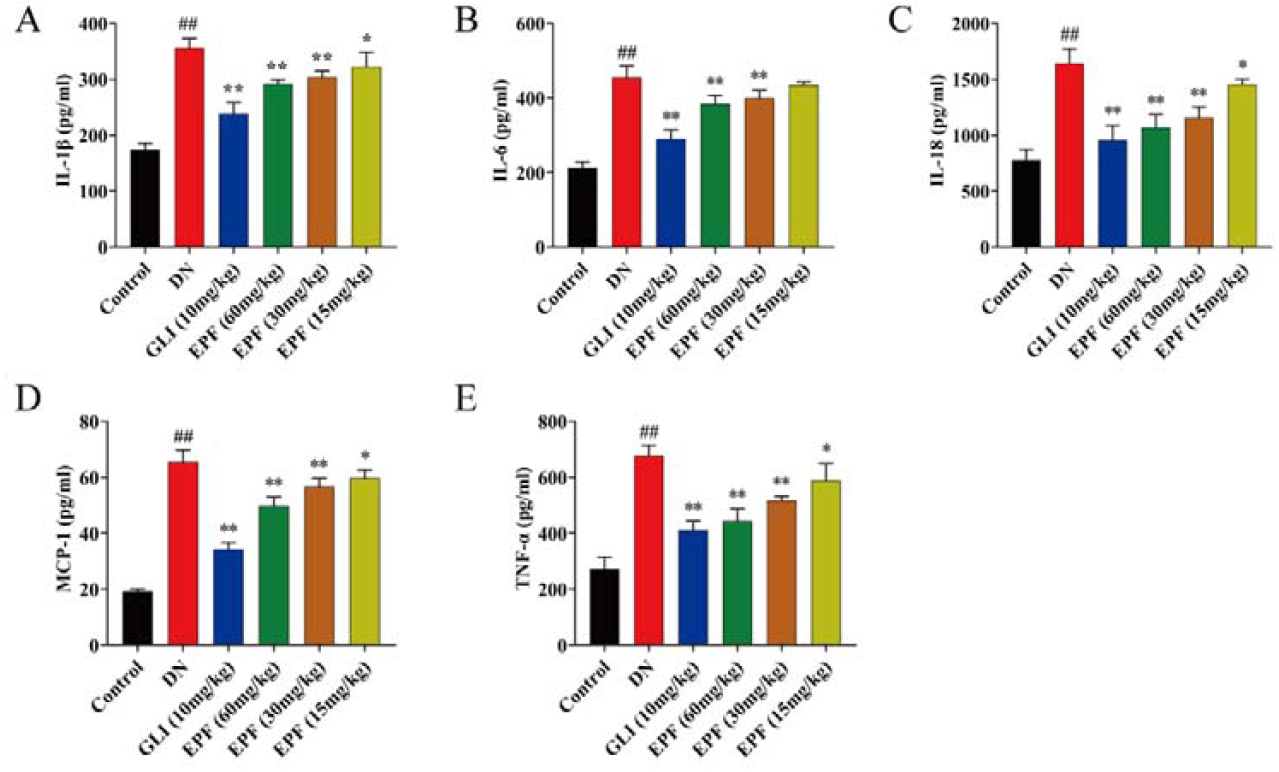
The effect of EPF on inflammatory factors in the serum of db/db mice (n=6). (A) Serum IL-1*β* level (B) Serum IL-6 level (C) Serum IL-18 level (D) Serum MCP-1 level (E) Serum TNF-*α* level. All data are expressed as mean ± S.E.M. ##P<0.01 compared with the control group; *P <0.05, **P<0.01 compared with the DN group

### 3.4 EPF Reduces Kidney Pathological Damage in db/db Diabetic Mice

The kidney tissue morphology indicated obvious kidney disease in the DN group, wit h lipomas around the kidneys and significantly larger kidney volume than what was observed in other groups. To explore EPF’s protective effect on diabetic kidney injury more directly, we used H&E and PAS staining on pathological sections of kidney tissues. The staining results showed that compared with the NC group, severe pathological damage was seen in the DN group. Part of the glomeruli and renal tubule s were completely infiltrated by inflammatory cells, and there was also glomerular hypertrophy, glomerular basement membrane thickening, and shearing of kidney tubule epithelial cells. After drug intervention (especially in the GLI group), pathological damage in EPF high-and medium-dose groups decreased significantly compared with that in the DN group. Although there was less improvement observed in the EPF low-dose group than in other treatment groups, greater improvement was found than that in the DN group (Fig. 4A-C)

**Figure 4.**
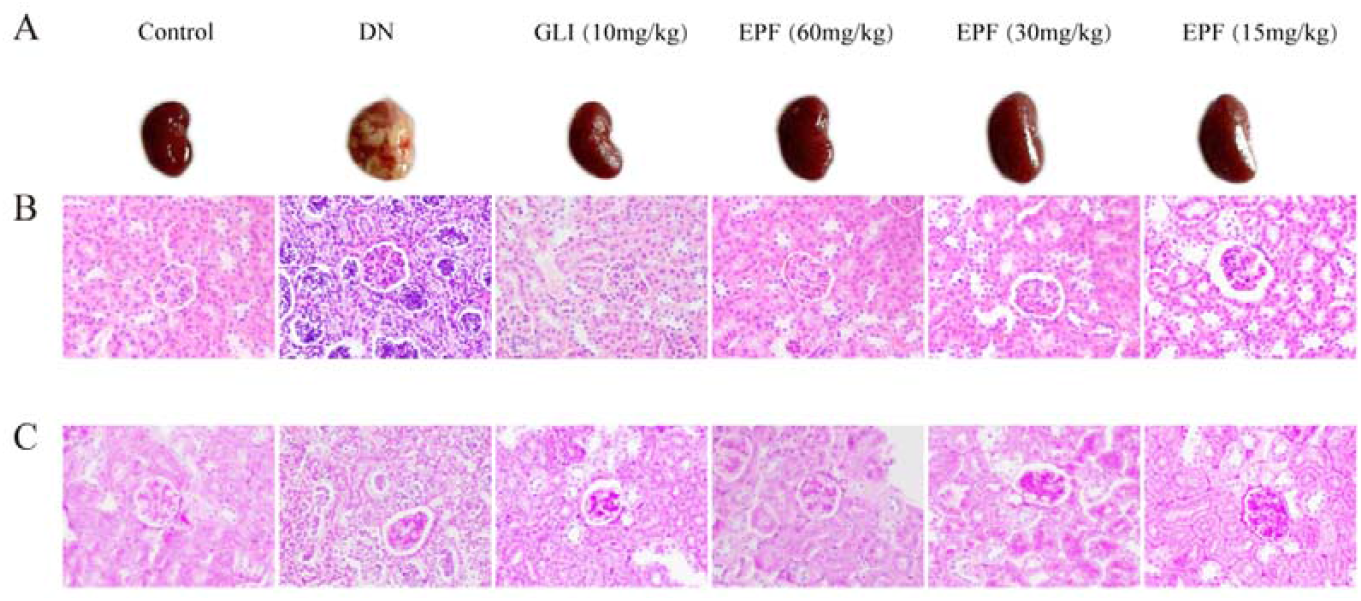
The effect of EPF on the pathological damage of kidney tissue in db/db mice. (A) Kid ney morphology (B) HE staining (400×) (C) PAS staining (400×).

To further clarify the effect of EPF on the signaling pathway, we used immunohisto chemistry (IHC) to measure the expression of TLR4, MyD88, NF-*κ*B, and MMP-9 factor s in the TLR4/MyD88/NF-*κ*B pathway in the mouse kidney interstitium. Compared with the negative control (NC) group, the expression of TLR4, MyD88, NF-*κ*B, and MMP-9 in renal tubular epithelial cells of diabetic nephropathy (DN) mice was enhanced. After EPF and GLI treatment, the expression of these factors all significantly decreased (Fig. 5A-B), which shows that EPF can significantly decrease kidney damage in db/db mice.

**Figure 5.**
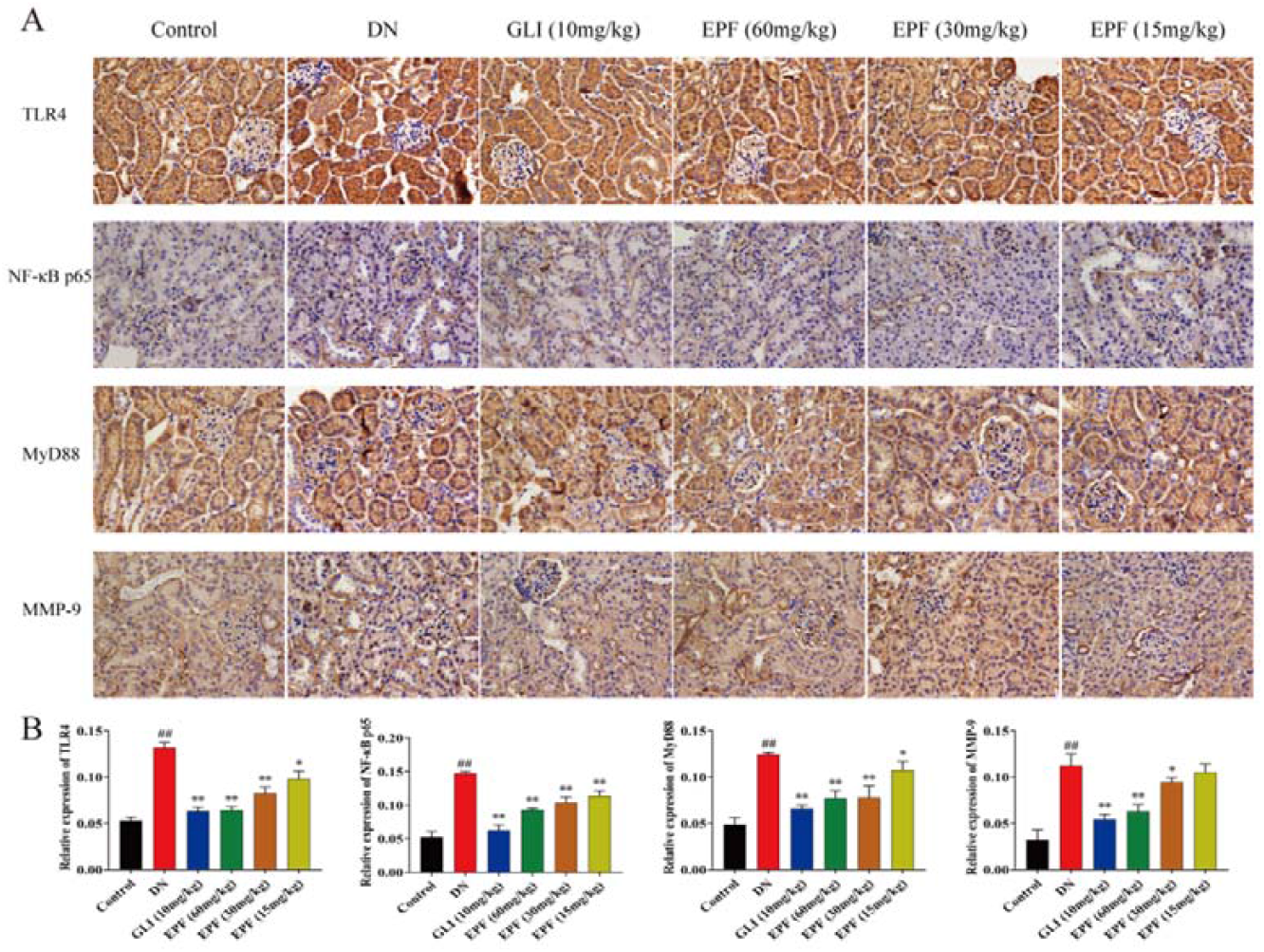
The effect of EPF on the levels of TLR4, p-NF-*κ*B p65, MyD88 and MMP-9 in the kidney of db/db mice. (A) Immunohistochemical map of TLR4, NF-*κ*B p65, MyD88 and MMP-9 (400×) (B) Imm unohistochemical optical density analysis of TLR4, NF-*κ*B p65, MyD88and MMP-9 proteins in kidne y tissue (n=3). All data are expressed as mean ± S.E.M. ##P<0.01 compared with the control group; *P<0.05, **P<0.01 compared with the DN group.

### 3.5 EPF Regulates the TLR4/MyD88/NF-*κ*B Signaling Pathway and Subsequently Improves Diabetic Nephropathy in db/db Mice

TLR4, NF-*κ*B p65, and MMP-9 are classic inflammation markers. To study whether EPF has a regulatory effect on the TLR4/MyD88/NF-kB signaling pathway, we measured the expression levels of TLR4, MyD88, NF-kB, and MMP-9 in mouse kidney tissue using WB and qPCR methods. The results showed that in the WB experiment, the expression of TLR4, MyD88, NF-*κ*B, and MMP-9 proteins in the GLI group and the high, medium, and low dose EPF groups was decreased to varying degrees compared with the DN group. Similarly, in the qPCR experiment, the mRNA expression levels of TLR4, MyD88, NF-*κ* B, and MMP-9 were reduced in the GLI group and the EPF group compared with the model group (Fig. 6A-C). This suggests that EPF reduces kidney inflammation caused by diabetic nephropathy by regulating the TLR4/MyD88/NF-*κ*B signaling pathway.

**Figure 6.**
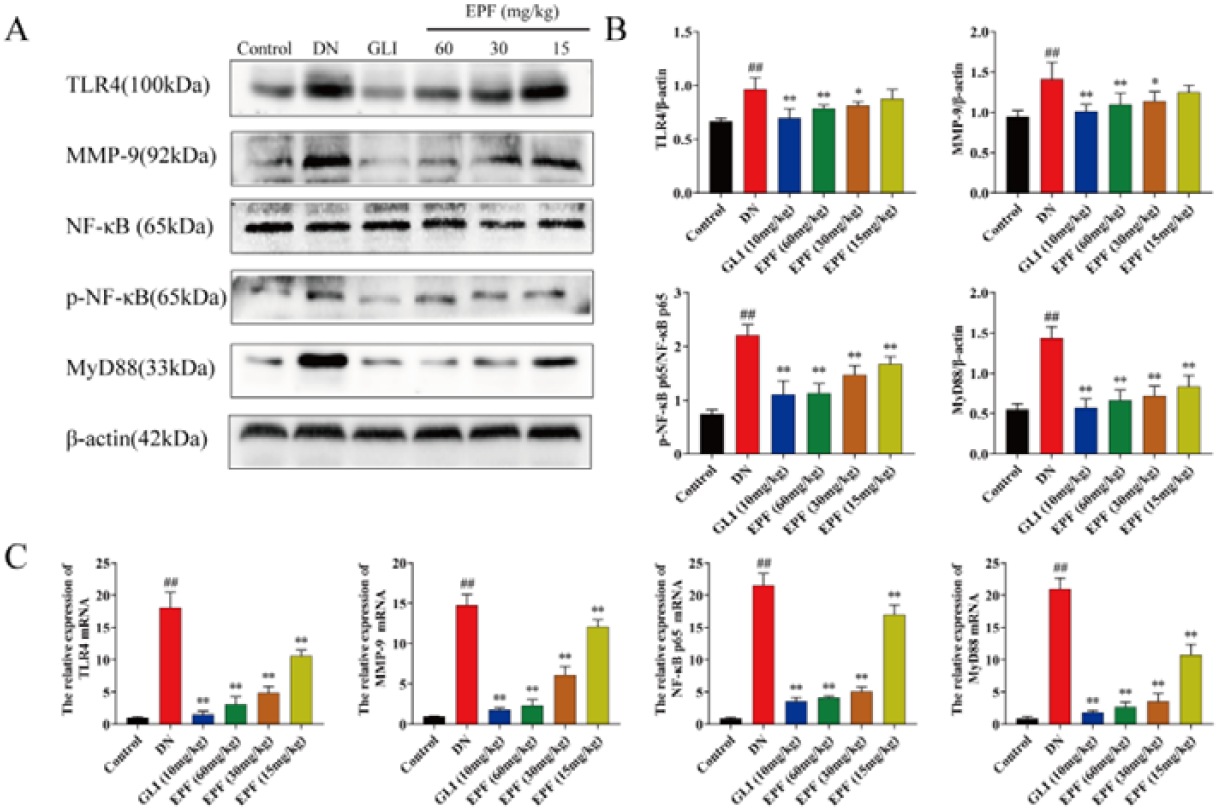
The effect of EPF on the TLR4/MyD88/NF-*κ*B signaling pathway in the kidney tissue of db/db mice. (A) Representative bands of TLR4,MMP-9, NF-*κ*B p65, p-NF-*κ*B p65 and MyD88 obtained by western blotting. (B) The relative expression of TLR4, MMP-9, p-NF-*κ*B p65/NF-*κ*B p65 and My D88 proteins (n=3). (C) The relative expression of TLR4, MMP-9, NF-*κ*B p65, and MyD88 mRNA (n=3). All data are expressed as mean ± S.E.M. ^##^P<0.01 compared with the control group; *P<0.05, **P<0.01 compared with the DN group.

### 3.6 EPF Has No Apparent Toxicity to KM Mice

After administering 300 times the dose of EPF to KM mice for 5 minutes, four mice showed reduced activity (4/6) without death; 27 minutes after administration, the activities of the mice normalized. No mice died during the 14-day observation period and there was no significant difference in body weight. Upon dissection, no abnormalities were visible to the naked eye in any organs and there was no significant difference in organ coefficients (Fig. 7A-D). There was also no significant diff erence in serum biochemical indicators when compared to untreated controls (Fig. 8 A-E). These data indicate that 600 times the dose of EPF had no obvious toxic effe ct on KM mice.

**Figure 7.**
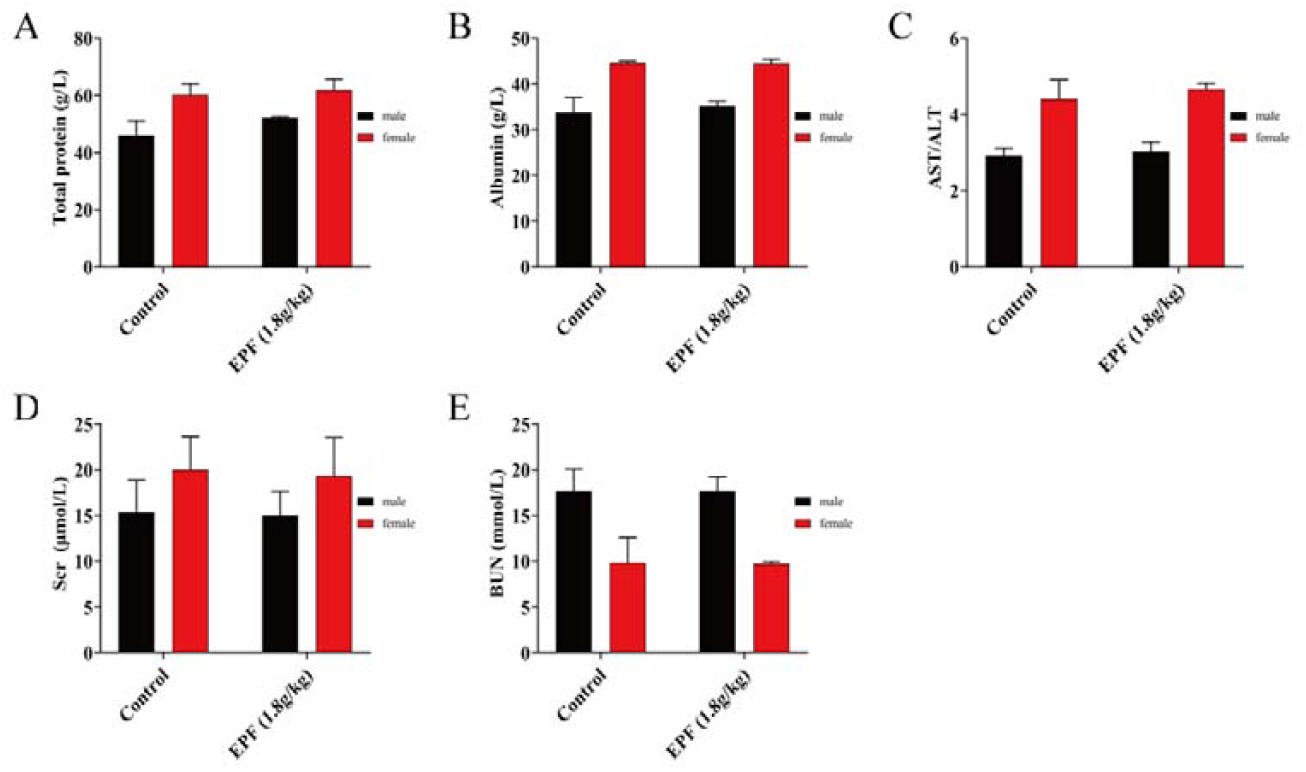
The effect of EPF on serum parameters of KM mice (n=3). (A) total protein (B) albumin (C) AST/ALT (D) Scr (E) BUN. All data are expressed as mean ±S.E.M.^##^P<0.01, ^#^P<0.05compared with the control group.

**Figure 8.**
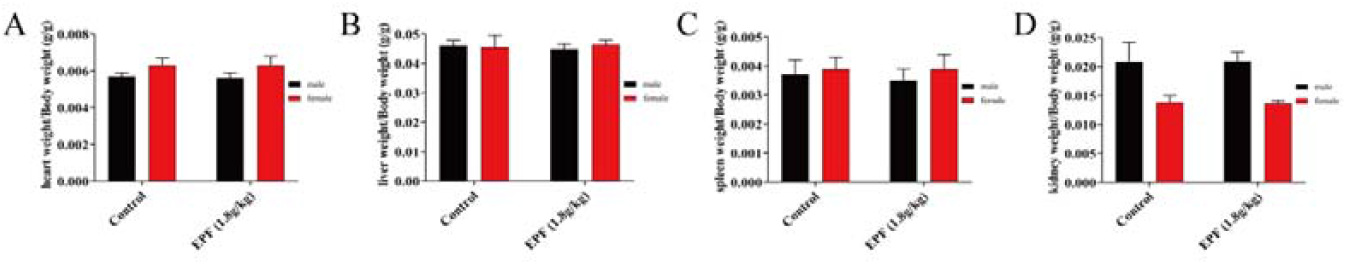
The effect of EPF on the organs/body weight of KM mice (n=3). (A) Heart weight/body weight (B) Liver weight/body weight (C) Spleen weight/body weight (D) Kidney weight/body weight. All data are expressed as mean ± S.E.M. ^##^P<0.01, ^#^P<0.05compared with the control group.

## 4 Discussion

It is estimated that by 2030, there will be more than 500 million diabetic patients worldwide, with DN affecting approximately half of them^17^. Because effective drugs for the prevention and treatment of DN are still insufficient at present, this disease and its complications pose a huge burden on human society. Severe metabolic abnormalities such as hyperglycemia, weight loss, and increased proteinuria are typical for diabetic patients. However, DN patients also experience renal tissue lesions such as glomerulosclerosis, glomerular basement membrane thickening, renal cell hypertrophy, and mesangial dilatation due to microvascular complications^18^. A s normal kidney function is gradually destroyed, countless DN patients eventually require repeated dialysis or kidney transplants^19^. Similarly, in our study, db/db mice exhibited symptoms such as hyperglycemia, weight loss, higher BUN and Scr lev els, and glomerulosclerosis. It is worth noting that EPF reduces blood glucose lev els and increases body weight in db/db mice while also reducing BUN and Scr levels

Inflammation plays a key role in the occurrence and development of DN. Toll-like receptors (TLR) can activate downstream inflammation signaling pathways^20^. As a uni que pattern recognition receptor, toll-like receptor 4 (TLR4) not only recognizes pathogen-related molecular patterns (PAMP) but also recognizes endogenous damage-related molecular patterns (DAMPs), including the production of lipopolysaccharides and fatty acids. During the occurrence and development of DN, high glucose and angiotensin II (AngII) significantly promote the expression and activation of TLR4^21^. A study^22^ showed that there is a high expression of TLR4 and other inflammatory mediators in DN rats. Similarly, in our study, the TLR4 level of the DN group was significantly higher than that of the NC group. However, after EPF treatment, the expression of TLR4 in mice was significantly reduced.

NF-*κ*B can be activated by TLR4, and the use of the TLR4/NF-*κ*B signaling pathway is a classic method to initiate internal inflammatory signal transduction^23^. Gener ally, activated TLR4 stimulates NF-*κ*B-related signaling pathways through MyD88-dependent pathways to promote the release of inflammatory cytokines and chemokines, such as MCP-1, IL-6, IL-8, IL-18, tumor necrosis factor-alpha (TNF-*α*), and matrix metalloproteinase-9 (MMP-9). MMP-9 is an important inflammatory marker of diabetic nephropathy and the most important enzyme to degrade type IV collagen. Under patho logical conditions, it plays an important role in proteinuria and the remodeling o f the glomerular basement membrane (GBM). Studies have shown that the concentration of MMP-9 increases in patients with type II diabetes, and its high expression precedes mALB, which plays a certain role in the occurrence and development of diabetic kidney damage^24^. In addition, there is clear evidence that the activation of N F-*κ*B is involved in the development of renal inflammation and fibrosis after the onset of DN^25^. He^25^ also found that DN in rats was alleviated by inhibiting the activation of NF-*κ*B and then downregulating the levels of other inflammatory factors.

In our study, we found that the expression of MyD88, NFr-*κ*B, and downstream inflammatory cytokines and chemokines in the kidney tissue of mice with diabetic nephropathy (DN) was significantly higher than in the control group. However, treatment with EPF significantly reduced the expression of these factors. This suggests that EPF can reduce kidney inflammation in diabetic nephropathy by inhibiting the TLR4/ MyD88/NF-*κ*B signaling pathway.

EPF is a medicine extracted from P. fallax, known for its anti-inflammatory and an ti-aging effects. The mechanism by which EPF functions in diabetic nephropathy is not yet clear. In our study, we observed severe kidney inflammation, tissue damage, and dysfunction in mice with diabetic nephropathy. Treatment with EPF significan tly improved these conditions. We believe that EPF inhibits the expression of inflammatory proteins in the TLR4/MyD88/NF-*κ*B signaling pathway and normalizes import ant renal function indicators.

Our acute toxicity experiments showed that oral EPF has low toxicity and is well-tolerated even at high doses. This suggests that EPF is a safe and effective anti-inflammatory drug that can mitigate diabetic damage to renal function through the T LR4/nuclear factor-*κ*B pathway. Further studies are needed to evaluate its clinical role and safety in humans.

## 5 Conclusion

We found that EPF reduced inflammation by inhibiting the secretion of inflammatory cytokines and effectively improved diabetic nephropathy in db/db mice by inactivating the TLR4/MyD88/NF-*κ*B pathway. This suggests that EPF has the potential to be an effective treatment for diabetic nephropathy.

## Author contributions

Qin Xu, Yukun Bao provided experimental ideas. Yukun Bao, Zeyue Wang raised animals. Yukun Bao, Zeyue Wang completed all the experiments. Yukun Bao, Qing Xu edi ted the graphics. Yukun Bao, Lixin Wang, Qin Xu wrote and revised the manuscript. Yukun Bao,Zeyue Wang confirm the authenticity of all the raw data. All authors rea d and approved the final manuscript.

